# New integrative vectors increase *Agrobacterium rhizogenes* transformation and help characterise roles for soybean *GmTML* gene family members

**DOI:** 10.1101/2024.07.25.605222

**Authors:** Huanan Su, Mengbai Zhang, Estelle B. Grundy, Brett J. Ferguson

## Abstract

Hairy-root transformation is widely used to generate transgenic plant roots for genetic functional characterisation studies. However, transformation efficiency can be limited, largely due to the use of binary vectors. Here, we report on the development of novel integrative vectors that significantly increase the transformation efficiency of hairy roots. This includes pHGUS7, for promoter::reporter visualisation studies, and pHOG13, for genetic insertion and overexpression studies. These vectors have been designed to simplify cloning workflows, enhance the selection of positively transformed *Agrobacterium* colonies, and increase the transformation efficiency and ease of selection of genetically modified hairy roots. To demonstrate the efficacy of the new vectors, Too Much Love (TML) encoding genes acting in the Autoregulation Of Nodulation (AON) pathway of soybean were investigated. Both constructs provided significantly higher transformation rates than the binary vector control, often resulting in >70% of the roots being transformed. Overexpression of each individual TML encoding gene (*GmTML1a*, *GmTML1b* and *GmTML2*) using pHOG13 resulted in a significant reduction in nodule number, demonstrating the role of all three in inhibiting nodule organogenesis. Moreover, reporter-fusions with the promoter of each TML encoding gene using pHGUS7 revealed that each exhibits a unique pattern of expression in nodules, with *GmTML1b* displaying considerably stronger expression than *GmTML1a* or *GmTML2*. Taken together, these results demonstrate the utility and efficiency of the new pHOG13 and pHGUS7 integrative vectors in hairy-root transformation, and improve our understanding of the critical *TML-*encoding genes in soybean nodulation control.

## INTRODUCTION

Transgenic hairy roots can be used to functionally characterise genes of interest in plant roots. The process is much faster than whole plant transformation and avoids the need for labour intensive tissue culture steps. Using the technique, a transgene driven by a tissue-specific or constitutive over-expressing promoter can be inserted, or a gene of interest can be knocked-down (e.g. RNAi and CRISPRi) or knocked-out (e.g. genome editing, such as CRISPR), or promoter::reporter fusions can be inserted to visualise where a gene is transcriptionally active.

The technique uses *Agrobacterium rhizogenes* (*Rhizobium rhizogenes*) bacteria to generate hairy roots on the host plant. The method has been well established in various legume species, including soybean (*Glycine max*; e.g. Kereszt et al., 2007; Lin et al., 2011; Zhang et al., 2021; Chu et al., 2022), wild soybean (*Glycine soja*; e.g. Niu et al., 2020), common bean (*Phaseolus vulgaris*; e.g. Estrada-Navarrete et al., 2007; Ferguson et al., 2014), pea (*Pisum sativum*; e.g. Li et al., 2023), *Lotus japonicus* (e.g. Stiller et al., 1997; Shimamura et al., 2007; Xu et al., 2018) and *Medicago truncatula* (e.g. Crane et al., 2006; Zhang et al., 2020). It has been instrumental in characterising the function of genes involved in the environmentally- and agriculturally-important legume-rhizobia symbiosis that leads to the development of root nodules, in which critical nitrogen fixation takes place (e.g. Ferguson et al., 2010; Roy et al., 2021). This includes key components of the Autoregulation of Nodulation (AON) process that provide the host plant with control over their extent of nodule formation (Ferguson et al., 2019).

When using the technique with crop legumes, many hairy roots can be generated per plant, with each root representing a novel transgenic event. However typically less than 20% of hairy roots formed are transformed. This is largely due to the use of binary vectors (Bahramnejad et al., 2019), where two separate insertion events are required to transform Transfer DNA (T-DNA) into the plant genome. This includes T-DNA from the endogenous Ri plasmid of the compatible *A. rhizogenes* strain that is required to induce hairy root formation, in addition to T-DNA from a binary vector generated *in vitro* that has been transformed into the *A. rhizogenes* bacteria to express the transgene cassette of interest.

Here, we report the development of novel integrative vectors that considerably enhance the transgenic efficiency of hairy roots. This includes pHOG13 for transgene insertion and over-expression studies, and pHGUS7 for promoter::reporter fusions studies using the GUS reporter gene. These vectors have been designed to integrate via homologous recombination into the Ri plasmid of the *A. rhizogenes* strain K599, which can be used to transform a range of plant species. As these vectors integrate into the Ri plasmid, only one insertion event is required to trigger both the development of hairy roots and insertion of the transgene cassette into the plant genome. To demonstrate the utility of these newly developed vectors, over-expression and promoter::GUS reporter fusion studies with soybean were performed using members of the *Too Much Love* (*TML*) gene family that are involved in the AON pathway (Zhang et al., 2021; Takahara et al., 2013; Tsikou et al., 2018). There are three *TML* genes in soybean: *GmTML1a, GmTML1b* and *GmTML2* (Zhang et al., 2021). Each gene significantly reduced nodulation when over-expressed using the pHOG13 vector. Moreover, their native promoters exhibited similar, but subtly distinct expression patterns using the pHGUS7 vector. These findings confirm the functionality of the new integrative vectors, and they support the likelihood of high functional redundancy amongst the three *GmTML* genes, with subtle yet discrete differences in expression patterns indicating the potential for additional and/or distinct biological roles.

## RESULTS

### Construction of *Agrobacterium rhizogenes* K599-compatible integrative vectors for hairy root transformation

Two integrative vectors, called pHOG13 (Fig. 1A) and pHGUS7 (Fig. 1B), were developed to improve the transgenic efficiency of *A. rhizogenes strain* K599 in insertion (including RNAi and genome editing) or promoter::reporter fusion studies, respectively. To maximise their effectiveness, only sequences deemed pivotal to their functionality were included. This involves a kanamycin resistance gene and associated regulatory components needed for antibiotic selection of the bacteria. In addition, a 606 bp mobility element (*RP4mob*) that allows the movement of plasmids from an intermediate *Escherichia coli* donor (i.e. HB101) to the final *Agrobacterium* recipient and an 809 bp homologous sequence (*K2*) required for sequence homology-directed recombination between the integrative vector and native Ri plasmid of *A. rhizogenes* K599 (pRi2659) were included (Figure 1C and 1D). Movement and recombination of the plasmids is achieved through tri-parental mating. An alternative 141 bp *bom* core mobility element, which is functional during conjugation from the pBR322 plasmid, was also tested but yielded a comparatively low integration efficiency (∼100 times lower, 1338±319 compared with 14 colonies, respectively).

**Fig. 1.**
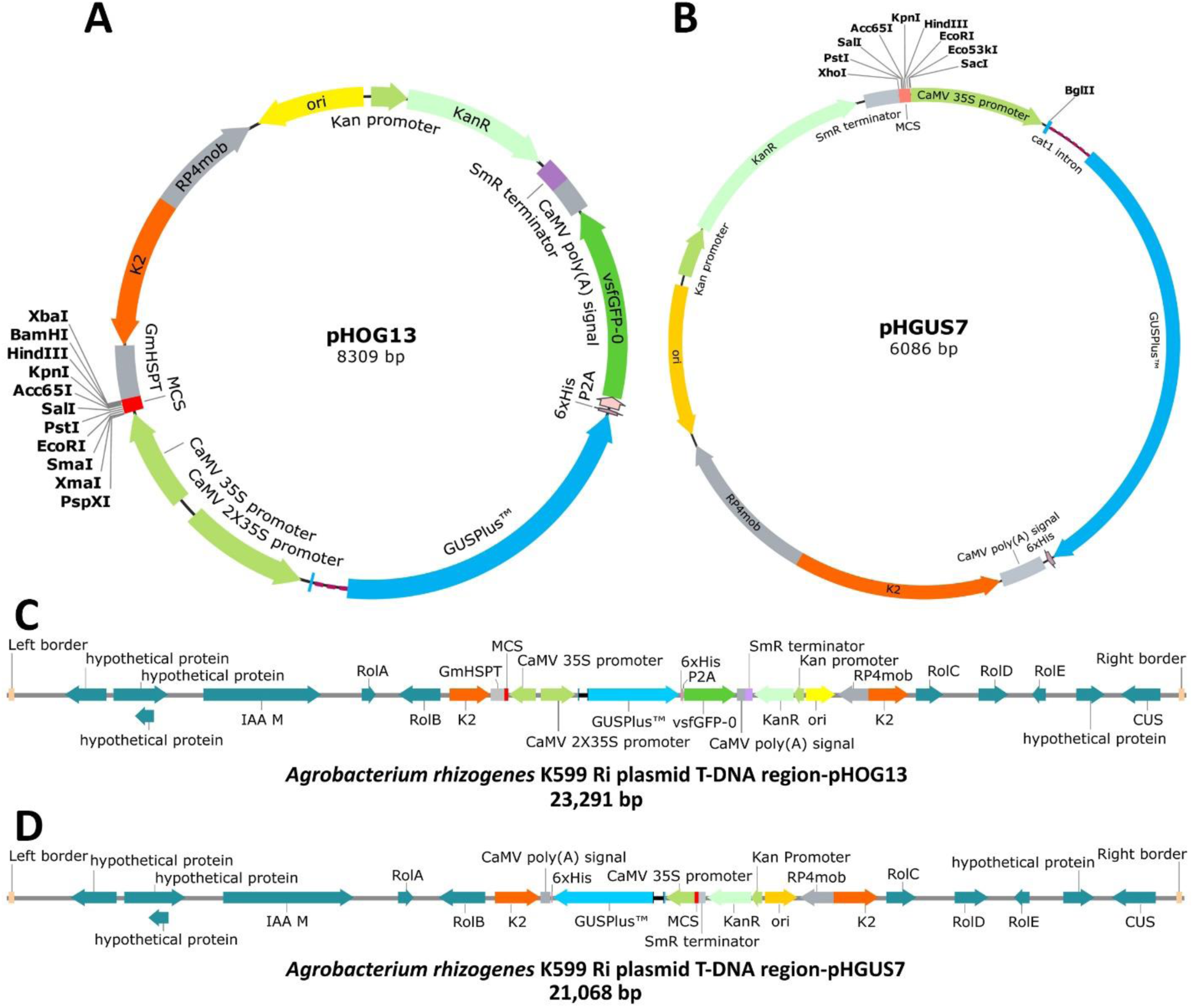
Sequences of pHOG13 and pHGUS7 integrative vectors for hairy root transformation with *Agrobacterium rhizogenes* strain K599. Vector maps of **A)** pHOG13 for transgene overexpression and **B)** pHGUS7 for *promoter::GUS* reporter analysis. Critical functional elements include: self-replication in *E. coli* (*Ori*, in yellow), tri-parental mating (*RP4mob*, in grey), integration into the *A. rhizogenes* K599 Ri plasmid (*K2* homologous region, in orange), selection of positive transformants in bacteria (e.g. kanamycin resistance gene, in light turquoise) and in plants (e.g. a *GUSPlus^TM^*/*vsfGFP-0* dual reporter gene, in blue and dark green), and transgene overexpression in hairy roots (e.g. MCS, in red). **C)** Gene configuration of the T-DNA segment of the *A. rhizogenes* K599 Ri plasmid. pHOG13 is integrated into the intergenic region between *RolB* and *RolC* and is flanked by a forward-repeated *K2* sequence within the T-DNA segment following tri-parental mating. **D)** pHGUS7 integrates into the same location of the Ri plasmid as pHOG13 due to the use of the same *K2* homologous sequence.

To enhance tri-parental mating with minimal false positives, antibiotic resistance testing was performed (Table S1). Interestingly, both *A. rhizogenes* K599 and K599-Rif (which is rifampicin resistant due to a naturally occurring mutation) were resistant to spectinomycin, making the *Spec^R^*gene unsuitable for our vector design. As a result, *Kan^R^* was chosen for the integrative vectors. Moreover, a high-copy-number origin of replication (*ori*) motif derived from the ColE1/pMB1/pBR322/pUC plasmid was incorporated into both integrative vectors, allowing for self-propagation in *E. coli* but not *Agrobacterium* cells. This was important to reduce background growth of rifampicin-resistant *A. rhizogenes* K599 that were transformed to contain the vectors without the occurrence of homologous recombination following tri-parental mating. The *pVS1 oriV* replication elements from *Pseudomonas* (widely used in binary vectors) were not included in these integrative vectors due to plasmid replication that would occur in K599 without an integration event, which causes false positives when screening with gene-specific primers on the destination K599 *Agrobacterium*.

Transgene overexpression using pHOG13 is enabled via inserting a coding sequence between a *CaMV 35S* promoter and *G. max* endogenous terminator at the multiple cloning site (MCS) which provides a range of restriction endonuclease recognition motifs. To facilitate screening in plants, an expression cassette having an enhanced *CaMV 35S* promoter-driven *GUSPlus* and *vsfGFP-0* dual reporter gene was included in pHOG13.

Visualisation studies using pHGUS7 are enabled via *promoter::GUS* fusion, where a *CAT1* intron-containing *GUSPlus* gene was cloned from pCAMBIA1305.1 and modified to be regulated by a *CaMV 35S* promoter and *CaMV poly(A) signal*, downstream of an MCS. To localise gene expression in roots, the *CaMV 35S* promoter in pHOG13 is simply replaced with a gene-specific promoter sequence of interest.

### Integrative vectors exhibit improved transformation efficiency compared with binary vectors

To establish the transformation efficiency of pHOG13 and pHGUS7, the integrative vectors were compared with the binary vector pCAMBIA1305.1 Δ2X35S for their ability to transform hairy-roots of soybean. Positively transformed roots were identified either through histochemical GUS-staining or/and detection of GFP fluorescence (Fig. 2A and 2B). Soybean plants inoculated with *A. rhizogenes* K599-Rif harbouring pHGUS7 or pCAMBIA1305.1 Δ2X35S produced a similar number of hairy roots, with an average of 17.6±1.55 and 18.2±1.03 per plant, respectively. However, the transformation efficiency of the integrative vector pHGUS7 was significantly higher at 57.21±2.85% with a 6.65±1.54% chimera rate, compared with the binary vector pCAMBIA1305.1 Δ2X35S at 26.88±1.36% with a 4.63±0.71% chimera rate (Fig. 2). The number of chimeric hairy roots was not significantly different between the two vectors (Fig. 2).

**Fig. 2.**
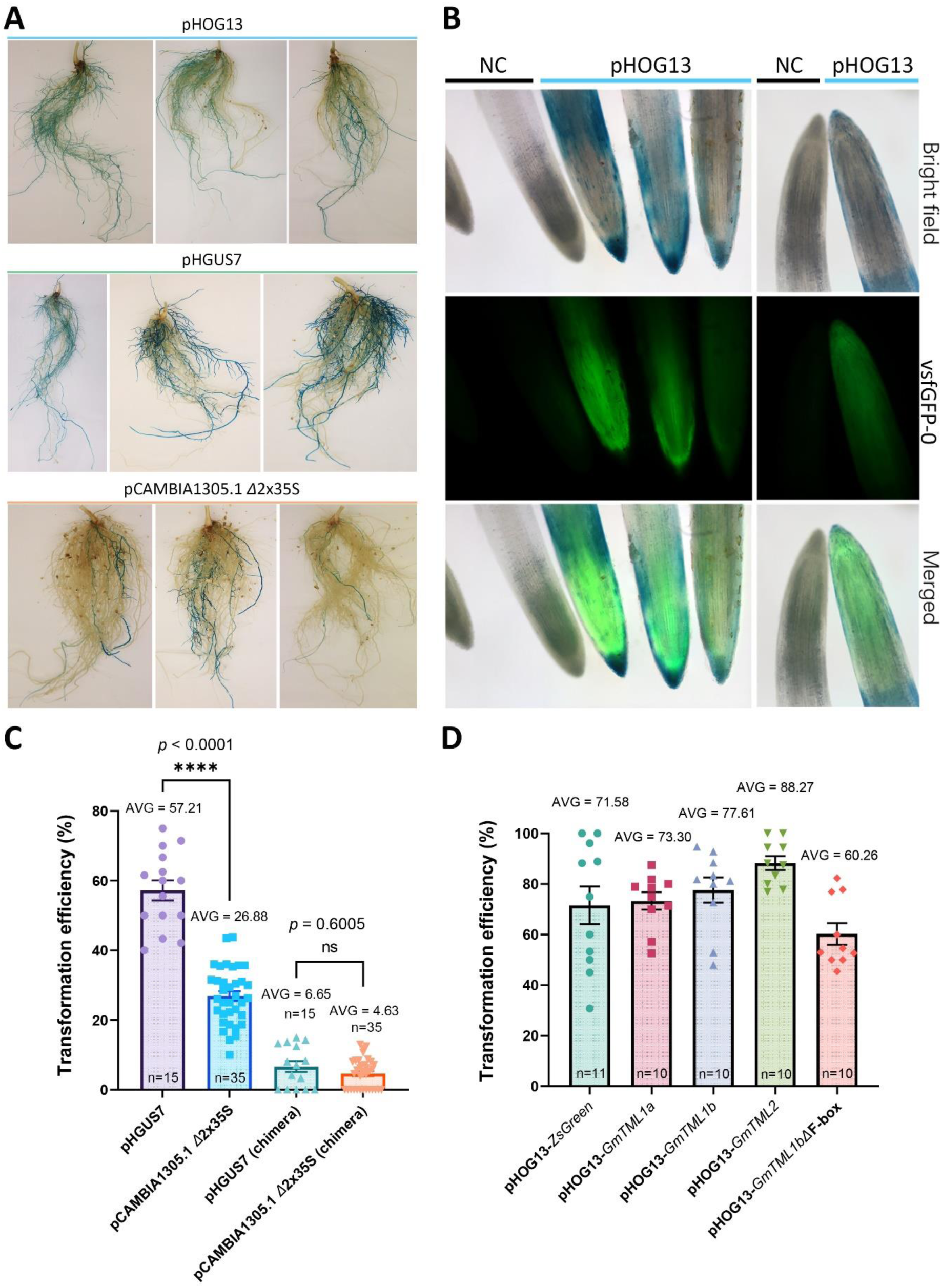
Use of the pHOG13 and pHGUS7 integrative vectors in *A. rhizogenes*-mediated hairy root transformation compared with the pCAMBIA1305.1 Δ2X35S binary vector. **A)** Hairy root systems of soybean plants transformed with empty pHOG13, pHGUS7 and pCAMBIA1305.1 Δ2x35S vectors following overnight histochemical GUS staining. **B)** Representative images of root tips of hairy roots transformed with an empty pHOG13 vector. Hairy roots were examined 2 h after staining using non-transformed soybean root tip as a negative control (NC). Green fluorescence was only detected in roots that were also positively stained for GUS. **C)** Efficiency of hairy root transformation using pHGUS7 and pCAMBIA1305.1 vectors quantified as the percentage of hairy roots emerging from the wound site that exhibited GUS activity. **D)** Efficiency of hairy root transformation amongst five different pHOG13-based overexpression constructs. Data are mean ± SEM with statistical significance indicated as asterisk.

To test the transformation efficiency of pHOG13, five over-expression constructs containing the coding sequences of *ZsGreen*, *GmTML1a*, *GmTML1b*, *GmTML2* and *GmTML1b* ΔF-box were generated and used for *A. rhizogenes* K559-mediated soybean transformation. Over-expression of the genes resulted in similar transformation efficiencies (71.58±7.45%, 73.3±3.47%, 77.61±4.97%, 88.27±2.76%, and 60.26±4.28% for *ZsGreen*, *GmTML1a*, *GmTML1b, GmTML2*, and *GmTML1b* ΔF-box, respectively) (Fig. 2D). This is similar to our findings using pHGUS7 (which shares the same *K2* homologous recombination sequence; Fig. 1) and is significantly higher than that observed with pCAMBIA1305.1 Δ2X35S (Fig. 2).

### Overexpression of *GmTML* genes reduces nodule numbers

To demonstrate utility of the pHOG13 integrative vector, the five constructs generated above were used to further understand the role of *TML* in the AON pathway of soybean. The pHOG13 vector containing only *ZsGreen* was used as the control. Full coding sequences of *GmTML1a*, *GmTML1b*, *GmTML2* were over-expressed and *GmTML1b* ΔF-box was used as a decoy (Feke et al., 2019). Due to a truncation of the functional F-box domain which has a structural role in bringing the target bound TML into the E3 ubiquitin ligase complex, the modified GmTML1b acts as a competitive binder. This protects a proportion of the TML protein targets from being ubiquitin-tagged for subsequent proteasomal degradation.

Transcript levels of *GmTML1a*, *GmTML1b* and *GmTML2* were significantly elevated in overexpressing hairy roots compared with control roots (Fig. 3A). *GmTML1a* overexpression exerted the strongest inhibition on nodule number, with *GmTML1b* or *GmTML2* overexpression also exhibiting a significant reduction in nodule numbers compared with control roots (Fig. 3B). In contrast, over-expressing the truncated *GmTML1b* increased nodule numbers. These findings demonstrate the reliability and robustness of the newly developed pHOG13 integrative vector, and further demonstrate the negative role that TML plays in the AON pathway of legume nodulation control.

**Fig. 3.**
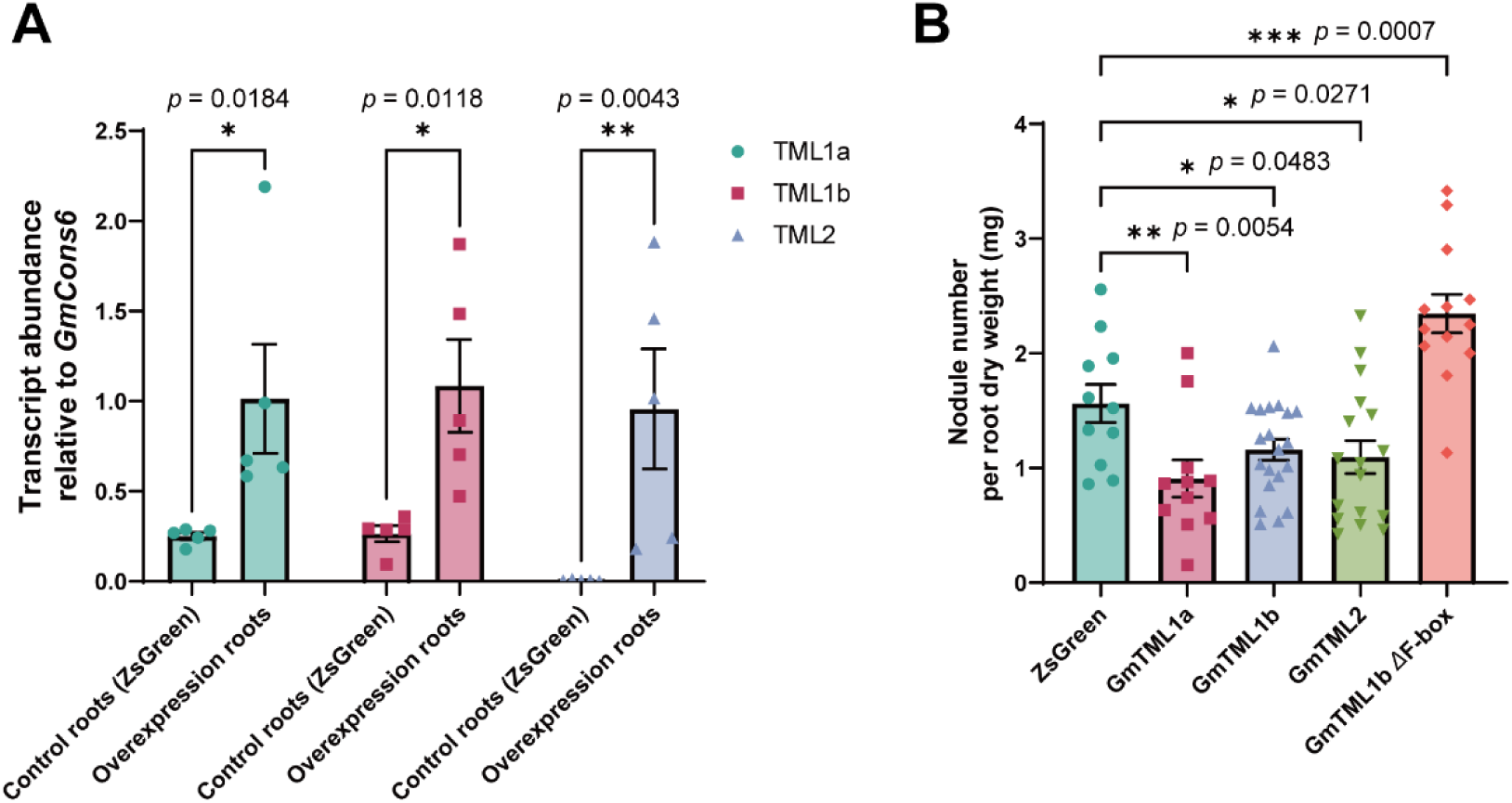
Over-expression of *GmTML* genes in hairy roots using the pHOG13 integrative vector and its effect on nodule development. **A)** Relative transcript abundance of *GmTML1a*, *GmTML1b* and *GmTML2* in individual hairy roots transformed with their respective pHOG13 over-expression construct. **B)** Nodule number phenotype in hairy roots transformed to overexpress the *ZsGreen* control gene, *GmTML1a*, *GmTML1b, GmTML2* and a truncated *GmTML1b*. Data are shown as mean ± SEM with asterisks indicating statistical significance. A) Two-way ANOVA; *n* = 5. B) One-way ANOVA with Fisher’s LSD test (n = 11-20).

### Localisation of *GmTML* expression reveals subtle differences in expression patterns

To demonstrate utility of the pHGUS7 vector for histochemical localisation of gene expression in roots, promoter::*GUS* fusion constructs of *GmTML1a*, *GmTML1b* and *GmTML2* were generated. Both *GmTML1a* and *GmTML1b* were transcriptionally induced in subepidermal cortical cells and pericycle cells during early rhizobia infection (Fig. 4A, E, H, L). Within young nodule meristems, their expression became more prominent in vigorously dividing cells (Fig. 4B, F, I, M). The transcriptional intensity of the gene homologs remained strong in nodule tissues where vasculature is formed, but became noticeably reduced in the cortical region that differentiates into the nodule infection zone (Fig. 4C, G, N). Both *GmTML1a* and *GmTML1b* genes were also expressed in nodule cortical and vascular tissues, with *GmTML1b* exhibiting slightly stronger intensity (Fig. 4D, G, K, O). As the nodule matured, the promoter activity of *GmTML1a* diminished and became confined to the nodule apex (Fig. 4D and 4G), whereas *GmTML1b* continued to be strongly expressed in nodule vasculature (Fig. 4K and 4O). Likewise, despite both being active in root vascular tissues, the expression of *GmTML1b* within the central stele was more persistent and broader (Fig. 4A-O).

**Fig. 4.**
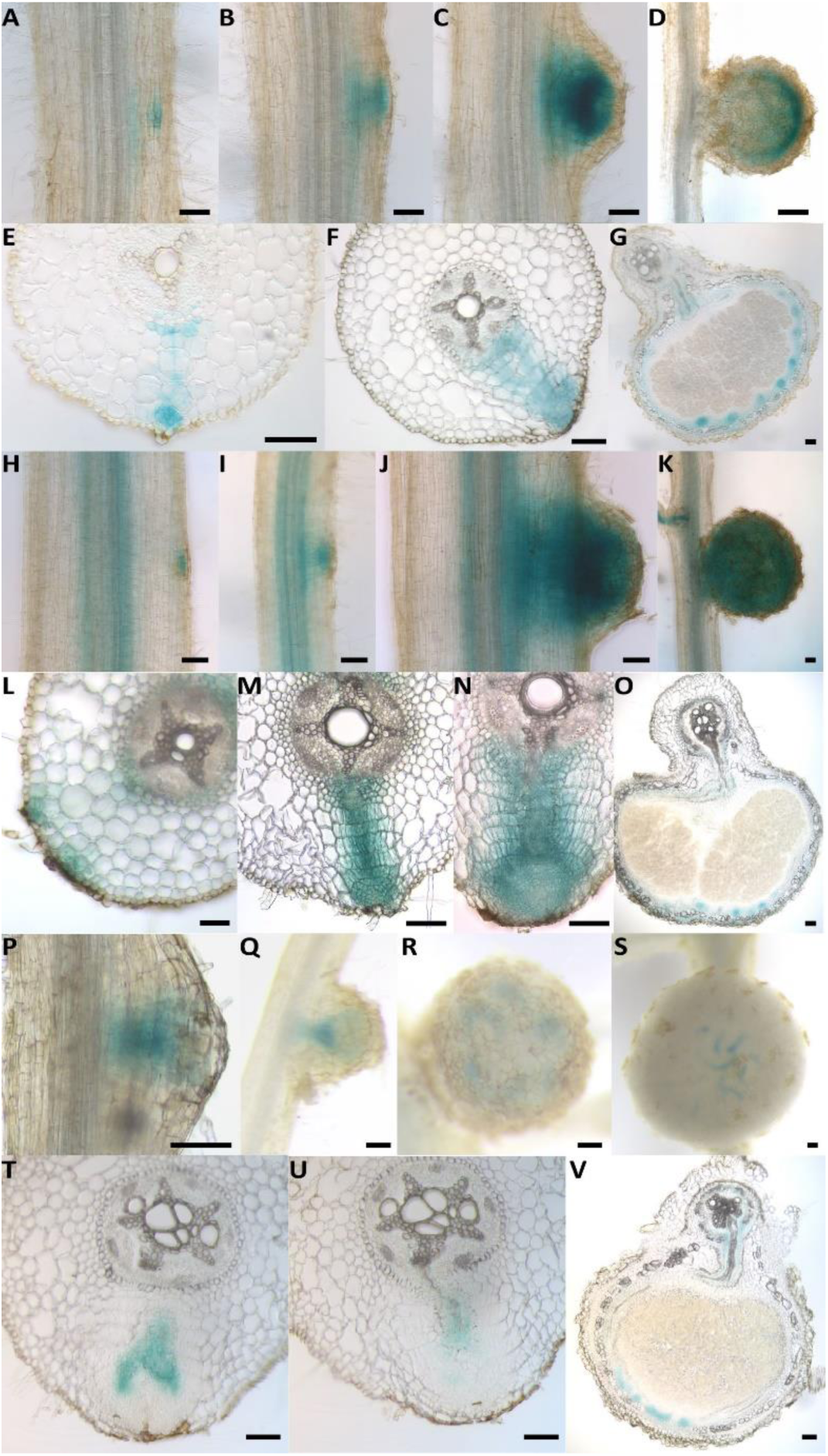
Histochemical localisation of the promoter activities of *GmTML1a*, *GmTML1b* and *GmTML2* obtained using pHGUS7 during nodule development. The expression patterns of *GmTML1a* (A-G), *GmTML1b* (H-O) and *GmTML2* (P-V) at various stages of nodule organogenesis were shown as whole-mount (A-D, H-K, P-S) and traverse-section (E-G, L-O, T-V) images. Scale bars represented 100 µm.

In contrast to the expression patterns of *GmTML1a* and *GmTML1b*, *GmTML2* was only weakly transcribed in root vasculature, and was not noticeably activated in the cortex until nodule vasculature developed (Fig. 4P, Q, T, U). At later stages of nodule organogenesis, *GmTML2* was only observed in newly formed vascular tissues (Fig. 4R, S, V). These findings suggest that considerable functional redundancy exists amongst *GmTML1a*, *GmTML1b* and *GmTML2*, and that *GmTML1* homologs may make a greater contribution to the AON nodulation control mechanisms based on their higher expression intensity.

## DISCUSSION

Hairy root transformation is a convenient and adaptable approach to functionally characterise genes expressed in roots. It is especially beneficial for species where whole-plant, stable transformation is challenging, resource-intensive and time-consuming. However, the standard use of binary vectors in hairy root transformation often necessitates the generation of numerous composite plants to produce sufficient quantities of positively transformed hairy roots. This is due to low transformation efficiency attributed to the need for two separate insertion events involving both the native Ri plasmid and introduced recombinant binary vector. This can be further exacerbated by the random and sometimes unfavourable integration of T-DNA into the plant genome.

In this study, we report on the development and utility of two novel integrative vectors that considerably enhance the efficiency of *A. rhizogenes* K599-mediated hairy root transformation. These integrative vectors, called pHOG13 and pHGUS7, resulted in a 30-100% (70% on average, n = 66) transformation efficiency of soybean hairy roots, compared with only 10-44% (27% on average, n = 35) using standard binary vectors. Incorporation of a *GUSPlus^TM^*/*vsfGFP-0* dual reporter in pHOG13 enabled the rapid selection of positively transformed hairy roots through histochemical staining of GUS activity or GFP visualisation. We noticed that some roots exhibited varying degrees of GUS staining, which may be attributed to the insertion event at different locations in the genome resulting in enhanced/repressed *GUS* gene expression, or different T-DNA copy numbers; however, no significant differences were observed between the integrative and binary vectors. Moreover, these reporters reveal the presence of chimeric roots, providing a visual means for selecting suitable transgenic roots for subsequent analyses.

To demonstrate the efficacy of the vectors, *TML* genes of soybean involved in the AON pathway of legume nodulation control were functionally characterised (Tsikou et al., 2018; Zhang et al., 2021). Soybean has three *TML* gene family members, called *GmTML1a, GmTML1b, and GmTML2* (Zhang et al., 2021). Individually overexpressing these three *GmTML* genes using pHOG13 resulted in significantly fewer nodules formed. Notably, *GmTML1a* overexpression resulted in the highest level of inhibition of the three genes, suggesting it may have the strongest role in controlling nodulation. GmTML1a and GmTML1b only differ in one residue within the F-box domain, suggesting the difference in their ability to inhibit nodule development is likely due to minor sequence variations within the Kelch repeat domain leading to altered target specificity or decreased binding affinity.

Our findings with *TML* are consistent with those reported in other legume species, which have varying copies of TML-encoding genes. Whereas soybean has three *TML*-encoding genes, *M. truncatula* has two, and *Phaseolus vulgaris* and *L. japonicus* and *P. sativum* have one (Takahara et al., 2013; Zhang et al., 2021; Chaulagain et al., 2023). Soybean underwent a whole genome duplication event ∼13 million years ago, resulting in a duplication of most genes (Schmutz et al., 2010). This is the reasoning for having two copies of *TML1* (*GmTML1a* and *GmTML1b*). In the case of *GmTML2*, its duplicate may have been lost over time, or it may have originated as a duplication from one of the *TML1* copies following the whole genome duplication and hence never had a duplicate (Zhang et al., 2021).

Our findings demonstrate that each of the soybean genes encoding TML reduce nodule numbers when overexpressed. This is consistent with reports based on *M. truncatula*, where overexpression resulted in fewer nodules (Gautrat et al., 2019). Moreover, loss of *LjTML* caused hyper-infection by rhizobia, followed by excessive nodule formation (Major et al., 2009; Takahara et al., 2013) and mutations in either *MtTML1* or *MtTML2* doubled the number of nodules formed compared to wild-type plants, and this increased to twenty-times more nodules in the double mutant (Chaulagain et al., 2023). Collectively, these findings highlight the role of TML as a negative regulator of nodule organogenesis in the AON pathway.

To establish expression patterns of the three *GmTML* genes, their promoters were fused directly upstream of the *GUS* reporter gene in pHGUS7. Similar, but distinct, patterns of expression were observed for each, and this varied with the stage of nodule development. Essentially, each of *GmTML1a, GmTML1b* and *GmTML2* exhibited expression from early through to late stages of nodule development, predominately located in developing vasculature and cortical cells and the nodule primordium. There was a conspicuous difference in the intensity of expression, with the promoter of *GmTML1b* exhibiting much stronger expression than that of *GmTML1a* or *GmTML2*. Consistent with our findings, *LjTML* is expressed in cortical and pericycle cells activated by rhizobia infection, as well as in nodule primordia (Takahara et al., 2013). Moreover, *MtTML1* (the ortholog to *GmTML1a* and *GmTML1b*) is induced in root vasculature shortly following rhizobia inoculation, and then is active in the nodule primordia and immature nodules (Chaulagain et al., 2023), reminiscent of the pattern reported here for *GmTML* genes and for *LjTML* (Takahara et al., 2013). In contrast, the expression of *MtTML2* is only localised in root vascular bundles during nodule organogenesis, indicating that *MtTML1* is functionally more specialised for AON (Chaulagain et al., 2023). Altogether, the expression patterns of *GmTML1a, GmTML1b* and *GmTML2*, and of *MtTML1* and *MtTML2*, suggest they may have subtly different roles in fine tuning root and nodule development, but overall may have a high level of functional redundancy.

Collectively, our findings demonstrate the effectiveness of pHOG13 and pHGUS7 as new tools for enhancing the transformation efficiency of *A. rhizogenes* K599 in hairy root studies. The K599 strain is effective in transforming several species, including the legumes *G. max* (soybean; Kereszt et al., 2007; Lin et al., 2011; Zhang et al., 2021; Chu et al., 2022), *G. soja* (wild soybean; Feng et al., 2022; Li et al., 2024)*, P. vulgaris* (common bean; Estrada-Navarrete et al., 2007; Ferguson et al., 2014)*, P. sativum* (pea; e.g. Li et al., 2023), *Lens culinaris* (lentil; Foti and Pavli, 2020), *Cicer arietinum* (chickpea; Aggarwal et al., 2018), *Vigna radiata* (mungbean; Cuong et al., 2023; Lin et al., 2022; Chen et al., 2022), *Vigna unguiculata* (cowpea; Ji et al., 2019), *Cajanus cajan* (pigeon pea; Meng et al., 2019), and non-legumes such as *Gossypium hirsutum* (cotton; Zhou et al., 2022), citrus species (Ma et al., 2022; Gong et al., 2024), *Cucumis sativus* (cucumber; Nguyen et al., 2022), *Cucurbita moschata* (pumpkin; Geng et al., 2022), etc. Some plant species having a high propensity for regeneration have also been used with K599 to generate stable culture-free lines, such as *Ipomoea batatas* (sweet potato; Cao et al., 2023, Mei et al., 2024), *Solanum tuberosum* (potato; Mei et al., 2024), *Taraxacum mongolicum* (dandelion; Cao et al., 2024), and succulent varieties (Lu et al., 2024). Expression of *WOX5* during the culture of callus and shoot induction can induce hairy root-to-shoot conversion after K599-mediated hairy root transformation in *Malus domestica* (apple tree; Liu et al., 2024) at a maximum rate of 20.6%. In future, pHOG13 and pHGUS7 can be modified further to contain mobility signals of different *A. rhizogenes* strains so they can be used to transform additional plant species.

## MATERIALS AND METHODS

### Plant materials and growth conditions

Wild-type soybean (*Glycine max* [L.] Merr. cv. Bragg) seeds were chlorine gas sterilised for 16 hours and then germinated in 4 L pots half-filled with fresh Grade 2 vermiculite using 28°C:26°C, 16 h:8 h, day:night conditions. Three-to-four day-old seedlings were subjected to hairy root transformation as outlined below. For nodulation studies, *Bradyrhizobium diazoefficiens* strain USDA110 was cultured in YMB medium at 28°C for three days and 150 mL of diluted rhizobia culture with OD600 at ∼0.1 was evenly applied to each pot.

### Construction of empty pHOG13 and pHGUS7 vectors

Vector construction of pHOG13 and pHGUS7 required a combination of strategies involving Gibson Assembly (NEB, Gibson Assembly Master Mix, E2611S), NEBuilder HiFi DNA Assembly (NEB, NEBuilder HiFi DNA Assembly Master Mix, E2621S), In-Fusion seamless cloning (TaKaRa, In-Fusion Snap Assembly EcoDry™ Master Mix, 638954), and Golden Gate Assembly (NEB, NEBridge Golden Gate Assembly Kit, E1601S). A workflow diagram for developing the vectors is provided in Figure S1, with a list of associated primers provided in Table S2. The vectors were optimised for 1) efficient integration into the endogenous Ri plasmid of *A. rhizogenes* K599, 2) avoidance of any unnecessary components or sequence to minimise the plasmid’s size, and 3) reliable selection of positively transformed bacterial colonies. In brief, the homologous recombination *K2* fragment was cloned from pRi2659/p15SRK2, the mobility component *RP4mob* was cloned from pK19mobSac8, the high-copy-number *ColE1/pMB1/pBR322/pUC* origin of replication (*ori*) was cloned from pUC19, the *SmR* terminator was cloned from p15SRK2, the *GmHSP* terminator was cloned from *G. max* (L.) Merr. cv. Bragg genomic DNA, the *CaMV 2x35S* promoter was cloned from pCAMBIA1302, and lastly the *Kan* promoter, *Kan^r^*, *CaMV poly(A) signal*, *GUSPlus*, and *CaMV 35S* promoter were cloned from pCAMBIA1305.1. The Multiple cloning site (MCS), *P2A*, and *vsfGFP-0* components were synthesised (Integrated DNA Technologies; IDT, Singapore or Genscript, China). Importantly, pHOG13 contains both a *CaMV 2X35S promoter*::*GUSPlus-P2A-vsfGFP-0*::*CaMV poly(A) signal* cassette for visualisation of positive transformants and a *CaMV 35S* promoter::MCS::*tGmHSP* for overexpression of genetic sequence(s) of interest. pHGUS7 does not contain these sequences, and instead has an MCS*-CaMV 35S* promoter::*GUSPlus*::*CaMV poly(A) signal* cassette. This enables the *CaMV 35S* promoter of pHGUS7 to be replaced with a different promoter sequence of interest, such as a promoter from a gene of interest, to visualise its expression pattern. All vector sequences were verified using Sanger and/or NGS (BatchSeq-plasmid) sequencing (Australian Genome Research Facility; AGRF).

### Development of over-expression and promoter::*GUS* fusion constructs

Full-length coding sequences of *GmTML1a*, *GmTML1b* and *GmTML2* were amplified from cDNA of root samples using PrimeSTAR Max DNA Polymerase (TaKaRa), using primers listed in Table S2. The resulting PCR products were column purified and treated with appropriate restriction enzymes to expose compatible sticky ends required for subsequent ligation into the digested pHOG13 over-expression vector. Sequences encoding a green fluorescent protein (*ZsGreen*) or a modified *GmTML1b* were synthesised by IDT (Singapore) and designed to be flanked by a *Hin*dIII and *Sal*I restriction site at either side in the pUCIDT-Amp cloning vector. Following digestion, the DNA fragments were gel purified and inserted into the pHOG13 vector at the MCS (Figure 1). NEB10β competent *E. coli* cells transformed with the ligation products were revitalised in SOC medium for 1.5 h in shakers and then cultured overnight on selective LB plates containing 50 µg/mL kanamycin at 37°C.

For promoter::reporter fusion studies, 4193 bp, 4245 bp, and 4154 bp sequences located directly upstream of the start codon of *GmTML1a, GmTML1b,* and *GmTML2*, respectively, were amplified from *G. max* (L.) Merr. cv. Bragg genomic DNA using primers listed in Table S2. The sequences were subsequently cloned into the pHGUS7 vector between the *Xho*I and *Bgl*II restriction sites in place of the *CaMV 35S* promoter, enabling gene-specific expression in hairy roots. All construct sequences were confirmed by Sanger sequencing at AGRF.

### Hairy root transformation

Integrative pHOG13 and pHGUS7 plasmids confirmed to have correct insert sequences were transformed into auxotrophic HB101 *E. coil* via heat shock. Positive HB101 clones containing the recombinant plasmids were then subjected to tri-parental mating with HB101 or DH5α *E. coli* containing the helper plasmid pRK2013 (kanamycin resistant) and *A. rhizogenes* K599 (rifampicin resistant), as described in Zhang et al. (2021). Notably, *Agrobacterium* minimal medium (MIN, Table S4) was used here to avoid *E. coli* overgrowth when selecting plasmid-integrated *Agrobacterium*. PCR was used to confirm successful transformation, with one set of primers to confirm the presence of the constructs and another to confirm homologous recombination between the *A. rhizogenes* Ri plasmids and the vectors of interest. Binary vectors were introduced into *A. rhizogenes* K599 cells via electroporation and positive clones were identified via liquid colony PCR. All primers are listed in Table S2.

Hairy root transformation was performed on three-and-a-half to four-day-old seedlings at 28°C:26°C day:night as described previously (Ferguson et al., 2014; Zhang et al., 2021; Chu et al., 2022). The seed coats were removed and the hypocotyls repeatedly (>50 times) stab-inoculated with the transformed *A. rhizogenes* K599 strains and mixed with 50 µM acetosyringone in sterile water. Cling wrap was used to cover the top of the pots to maintain high humidity and prevent the wound site from drying out. The pots were then maintained at 26°C:24°C day:night temperature to encourage *Agrobacterium* infection. After three days, the cling wrap was removed and pre-wetted Grade 2 vermiculite was added to the pots to cover the wound sites with the shoot exposed. This helps prevent the wound site from drying out to encourage maximum hairy root formation. Twice a week, 250 mL of a modified B&D solution (Broughton and Dilworth, 1971) supplemented with 2 mM KNO_3_ (Table S3) was provided to each pot. After roughly three weeks, the hairy roots had emerged and elongated to a stage where they could support the plant. At this stage, the original root system was excised to encourage growth and extension of the hairy roots, and the seedlings were transferred to new pots filled with fresh Grade 2 vermiculite and B&D nutrient solution supplemented with 2 mM KNO_3_. Two days after transplanting, the composite plants were inoculated with OD600 ≈ 0.1 *B. diazoefficiens* strain USDA110 to induce nodulation. The plants were then maintained at 28°C:26°C day:night and harvested 14 days post transplantation for assessing the extent of nodulation or for histochemical GUS staining.

### Histochemical staining, tissue sectioning and microscopy

For histochemical GUS staining, harvested plant tissues were vacuum infiltrated and incubated in a GUS reaction solution (100 mg/L X-Gluc, 0.516 mM potassium ferricyanide, 0.57 mM potassium ferrocyanide, and 5 mM EDTA in 1×PBS, pH 7.0) at 37°C for 6-12 h or 2 h (for checking both GUS and the vsfGFP-0 signal) after overnight fixation in 0.5% paraformaldehyde (prepared in 1×PBS, pH 7.0) (Jefferson et al., 1987). Cross-sections of hairy roots were achieved by embedding short root segments in agarose blocks (4-6%), followed by tissue sectioning using a vibratome VT1200S (Leica, Wetzlar, Germany) with 0.4 mm/s speed, 0.7 mm amplitude, and 60 µm thickness settings. Bright-field microscopy of whole or sectioned tissues was photographed using Nikon (Tokyo, Japan) CCD cameras (Nikon DS-Fi2 or Nikon DS-Ri1) connected to either a stereomicroscope (Nikon C-PS) or standard optical microscope (Nikon Eclipse E600).

### RNA extraction, cDNA synthesis and RT-qPCR analysis

RNA was isolated from plant tissues using a Maxwell RSC Instrument and Maxwell RSC Plant RNA Kit (Promega, Madison, WI, USA) following the manufacturer’s instructions. cDNA was then synthesised using 1 µg of total RNA in 20 μL reactions using SuperScript IV reverse transcriptase (Invitrogen, Waltham, MA, USA). Reverse transcription (RT)-quantitative real-time PCR (qPCR) was then performed using a CFX384 real-time system (Bio-Rad, Hercules, CA, USA) with Bio-Rad CFX Maestro 1.1 using GoTaq qPCR master mix (Promega). Relative expression was normalised against the *GmCons6* reference gene (Libault et al., 2008). Gene-specific primers are listed in Table S2.

### Statistical analysis

Statistical analyses were performed using GraphPad PRISM software version 9.1.2 (GraphPad Software, San Diego, CA, USA). Data were analysed using Student’s t-tests, one-way ANOVA, or Two-way ANOVA, with statistically significant differences labelled with asterisks (*, P < 0.05; **, P < 0.01; ***, P < 0.001 and ****, P < 0.0001).

## ACKNOWLEDGMENTS

This work was funded by an Australian Research Council Discovery Project to BJF (DP190102996). EBG was the recipient of an Australian Government Research Training Program (RTP) Scholarship. We would like to thank Dr Xitong Chu for assisting with the research, Dr Attila Kereszt for his contribution to developing the p15SRK2 vector and K599-Rif, and Dongxue Li for contributing to the construction of the pCAMBIA1305.1 Δ2X35S vector.

## CONFLICT OF INTEREST

The authors declare no competing interests.

## AUTHOR CONTRIBUTIONS

Huanan Su and Brett J. Ferguson: Designed the project. Huanan Su: Designed and assembled the new integrative vectors. Huanan Su, Mengbai Zhang and Estelle B. Grundy: Performed the experiments. Huanan Su, Mengbai Zhang and Brett J. Ferguson: Analysed the data. Huanan Su, Mengbai Zhang and Brett J. Ferguson: Wrote the manuscript. All authors helped edit the manuscript.

---

**Table S1.**
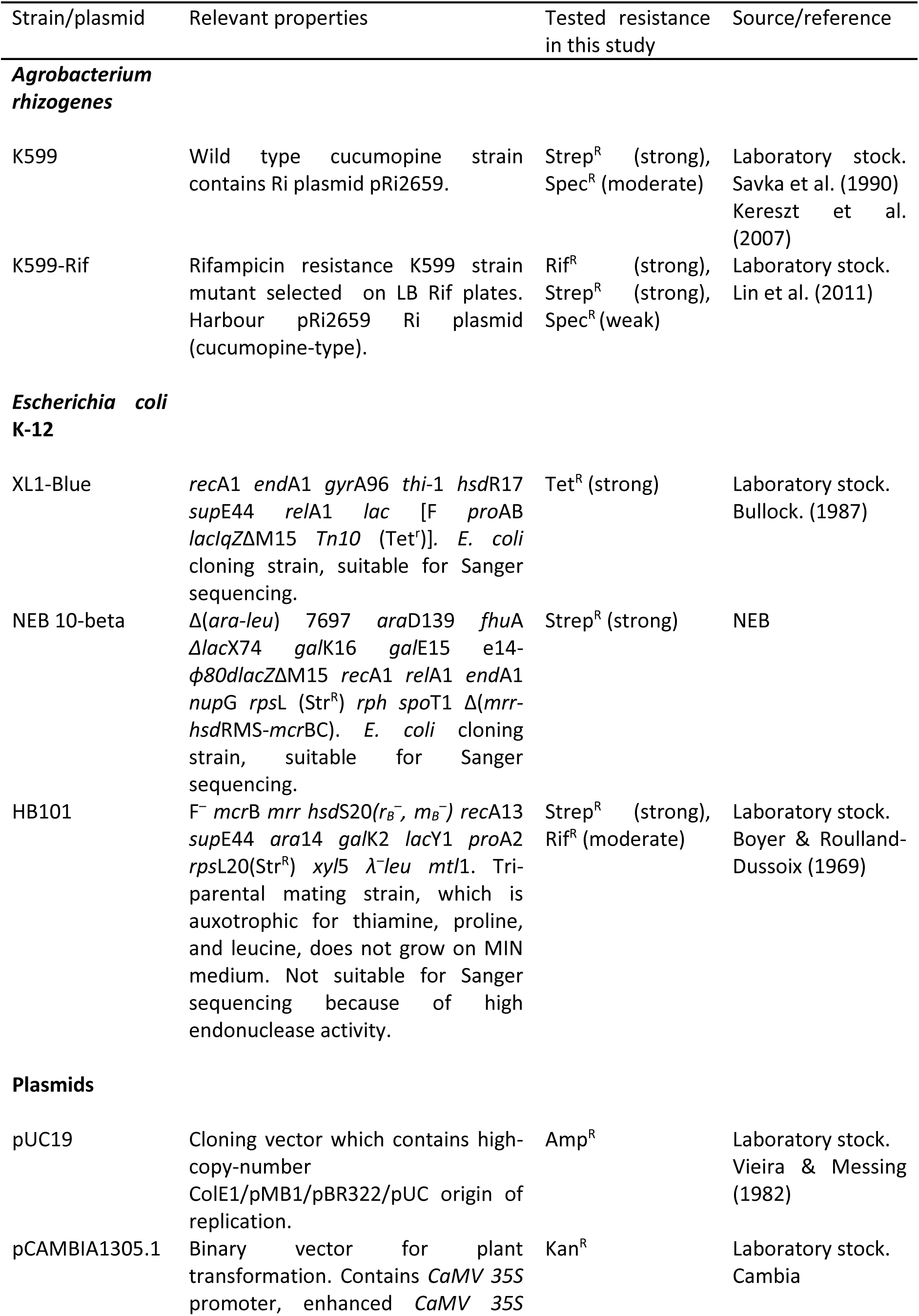
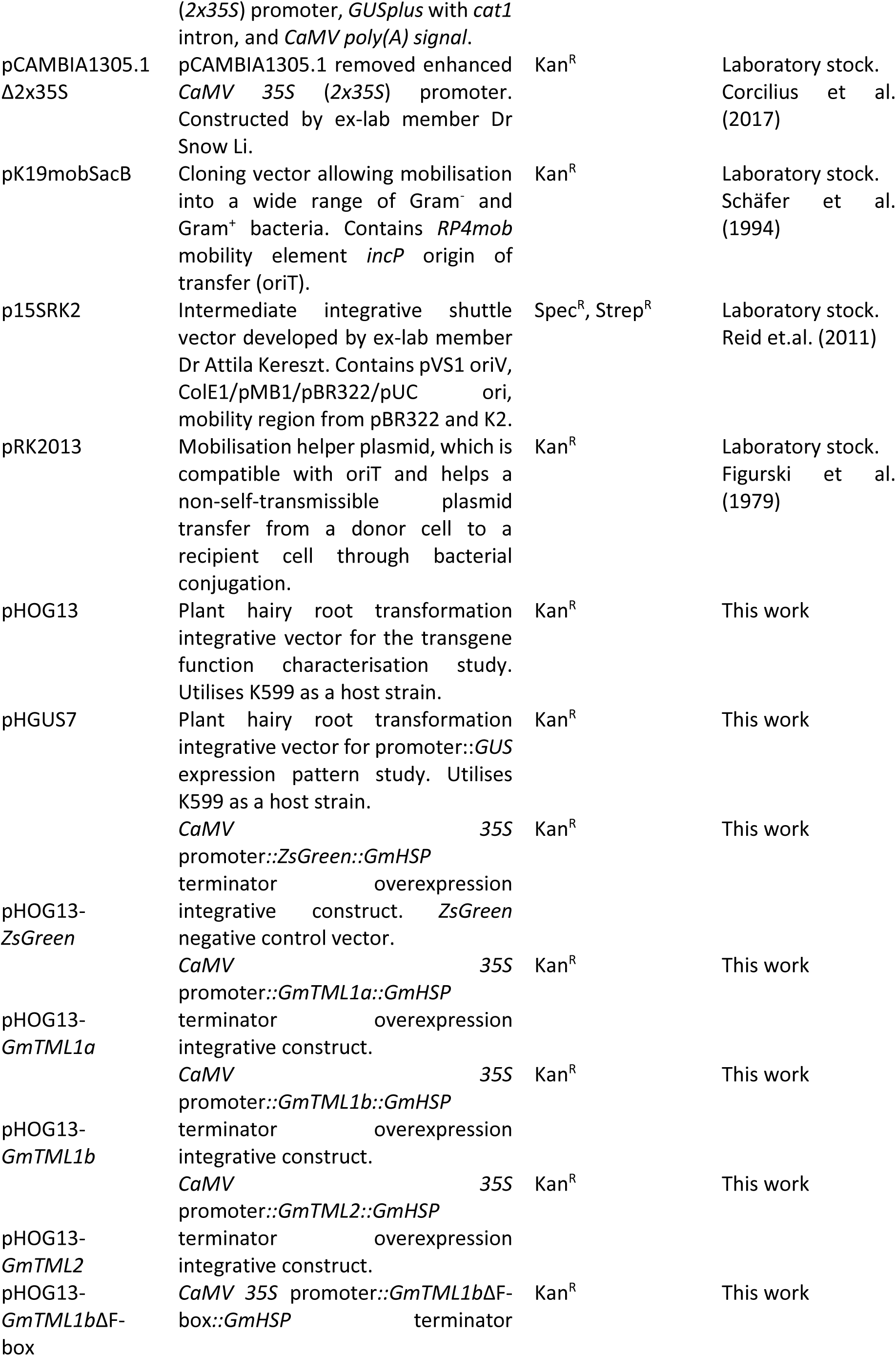
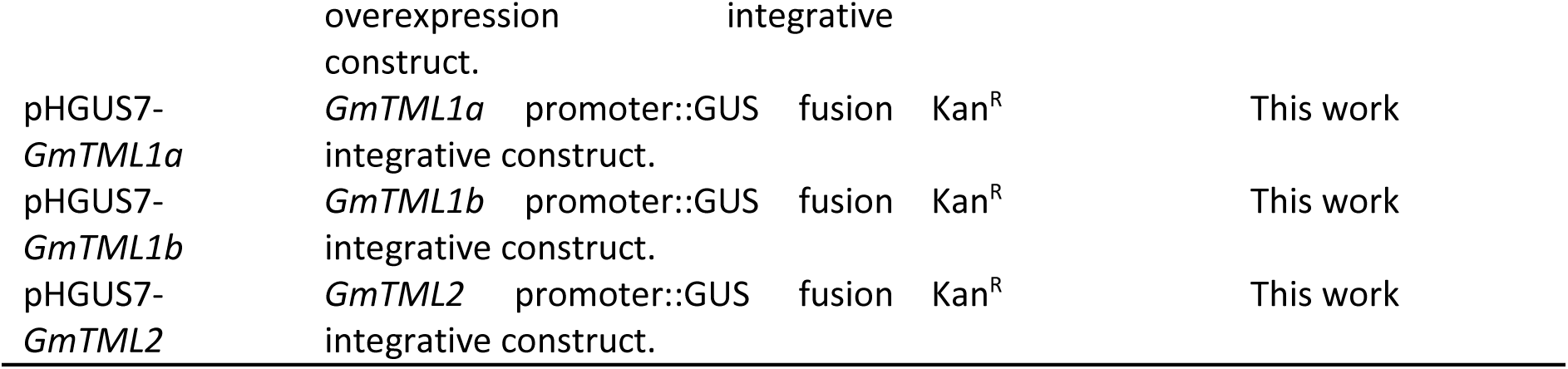
Bacterial strains and plasmids used in this study and their antibiotic resistance.

**Table S2.**
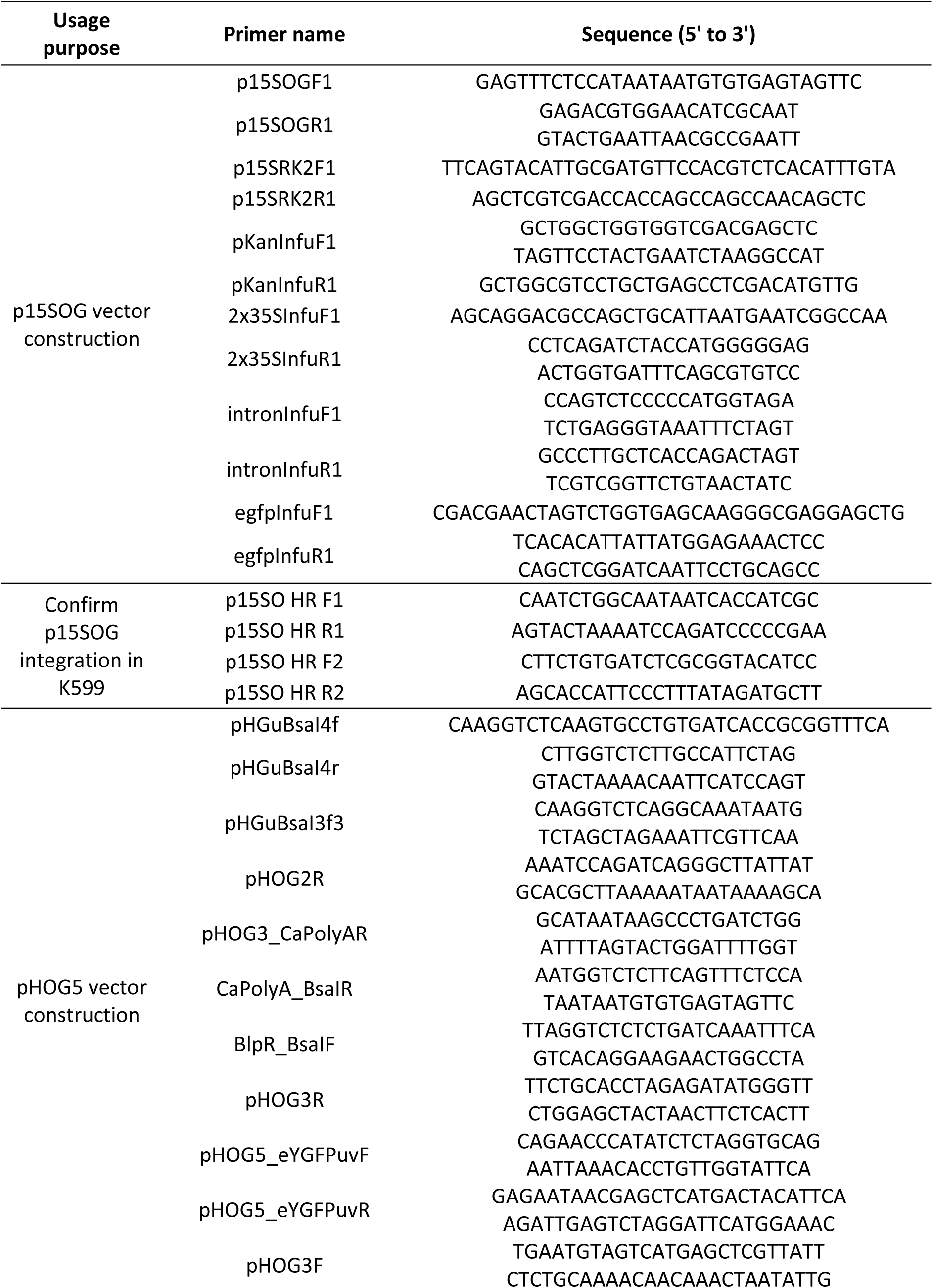
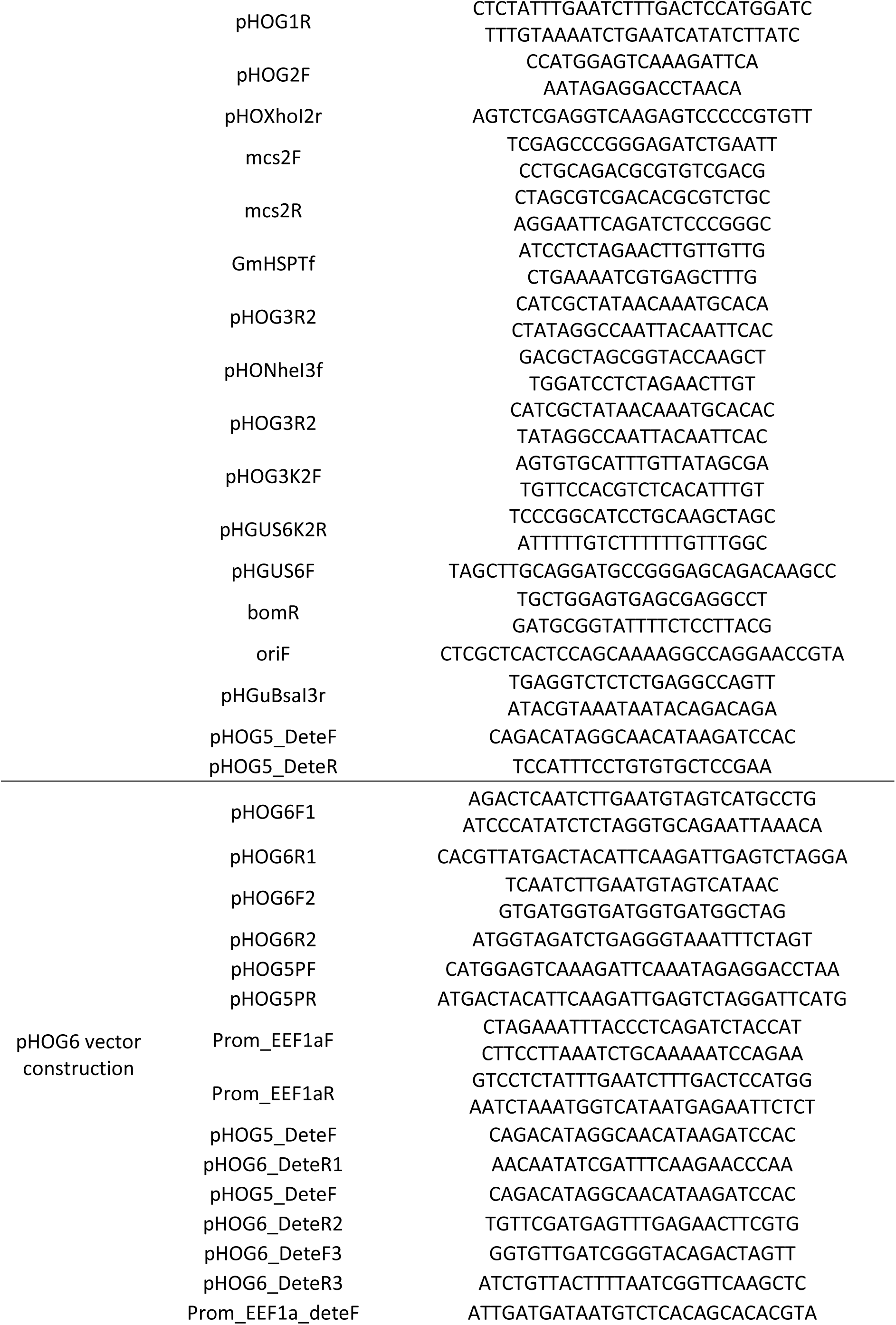
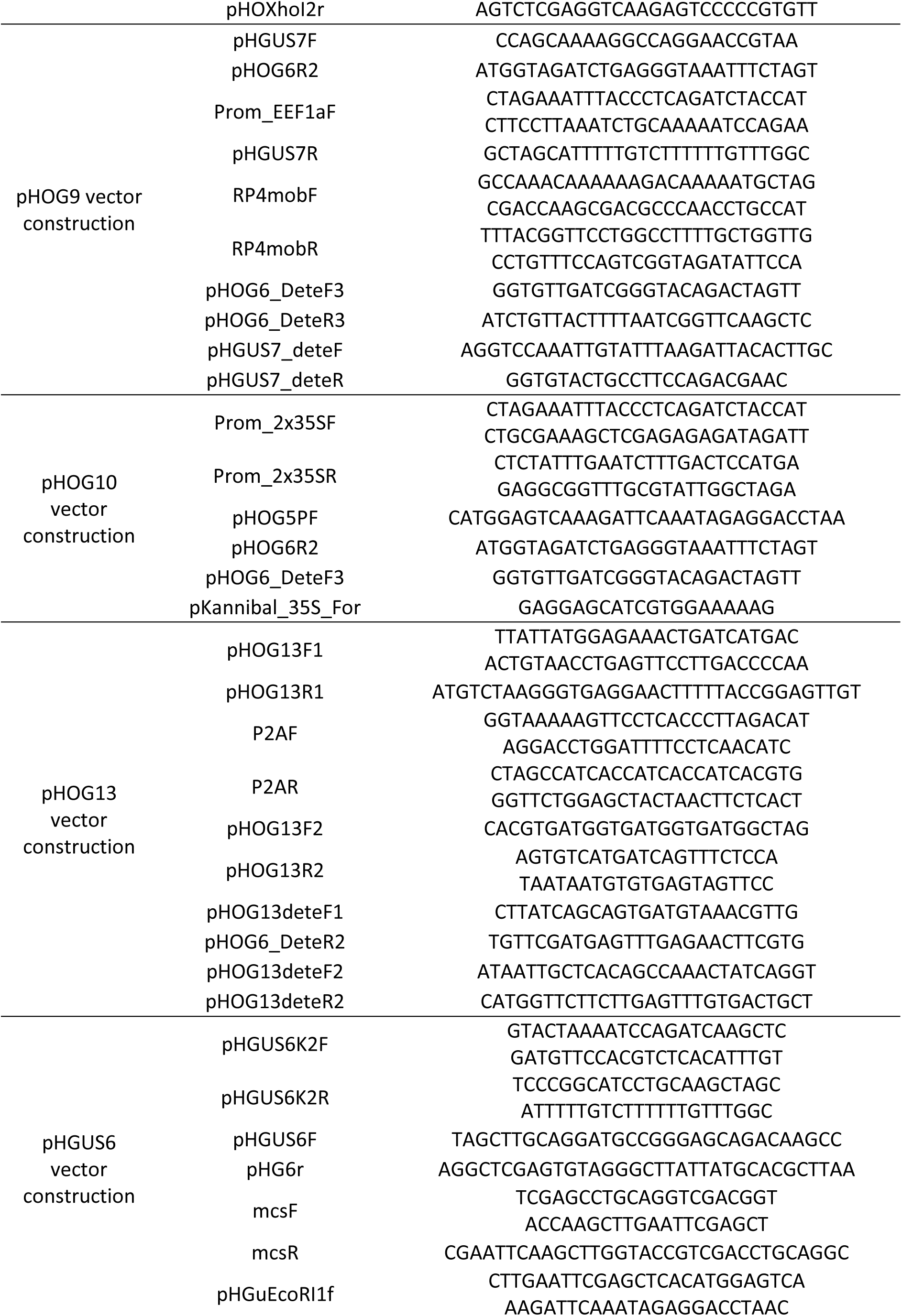
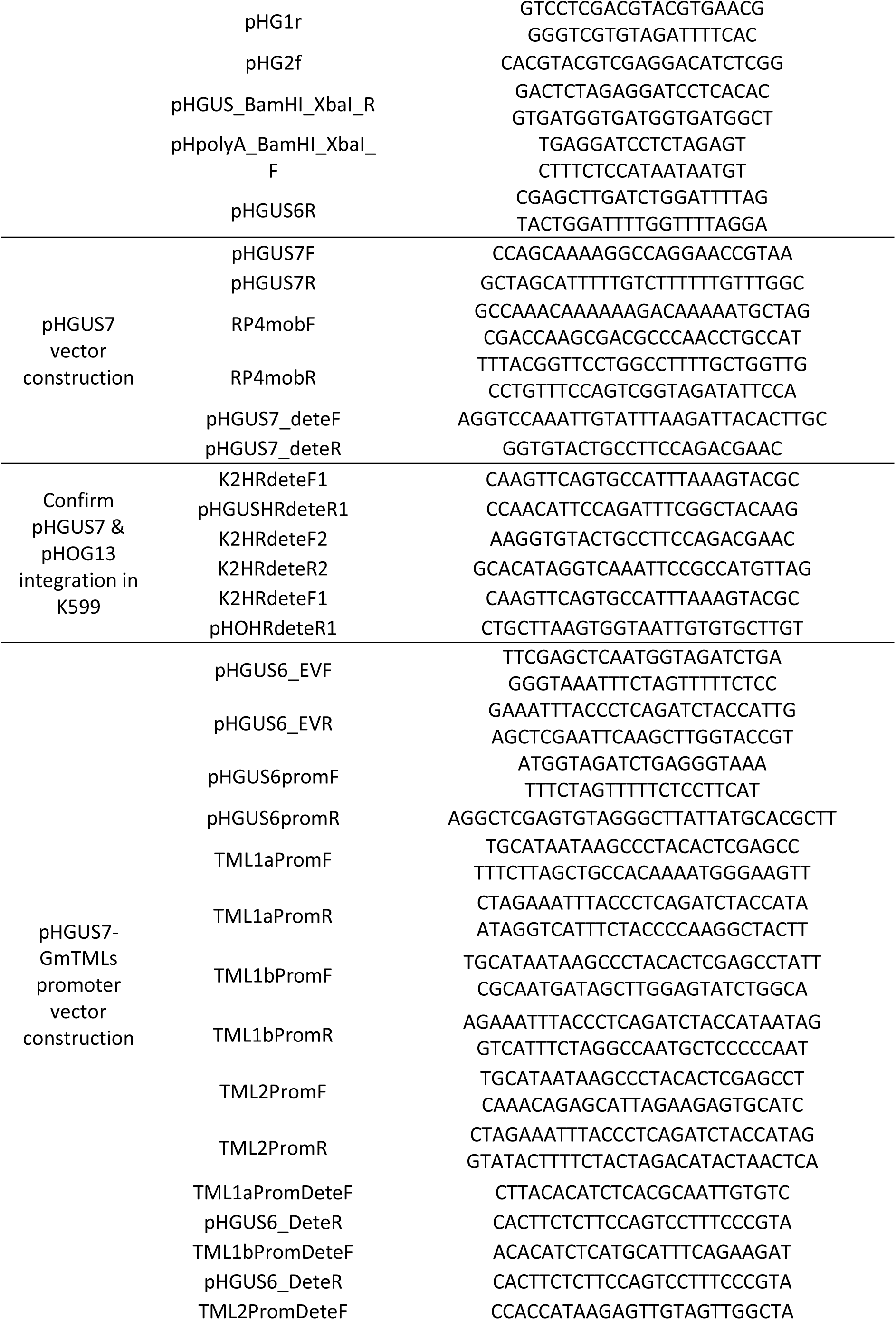
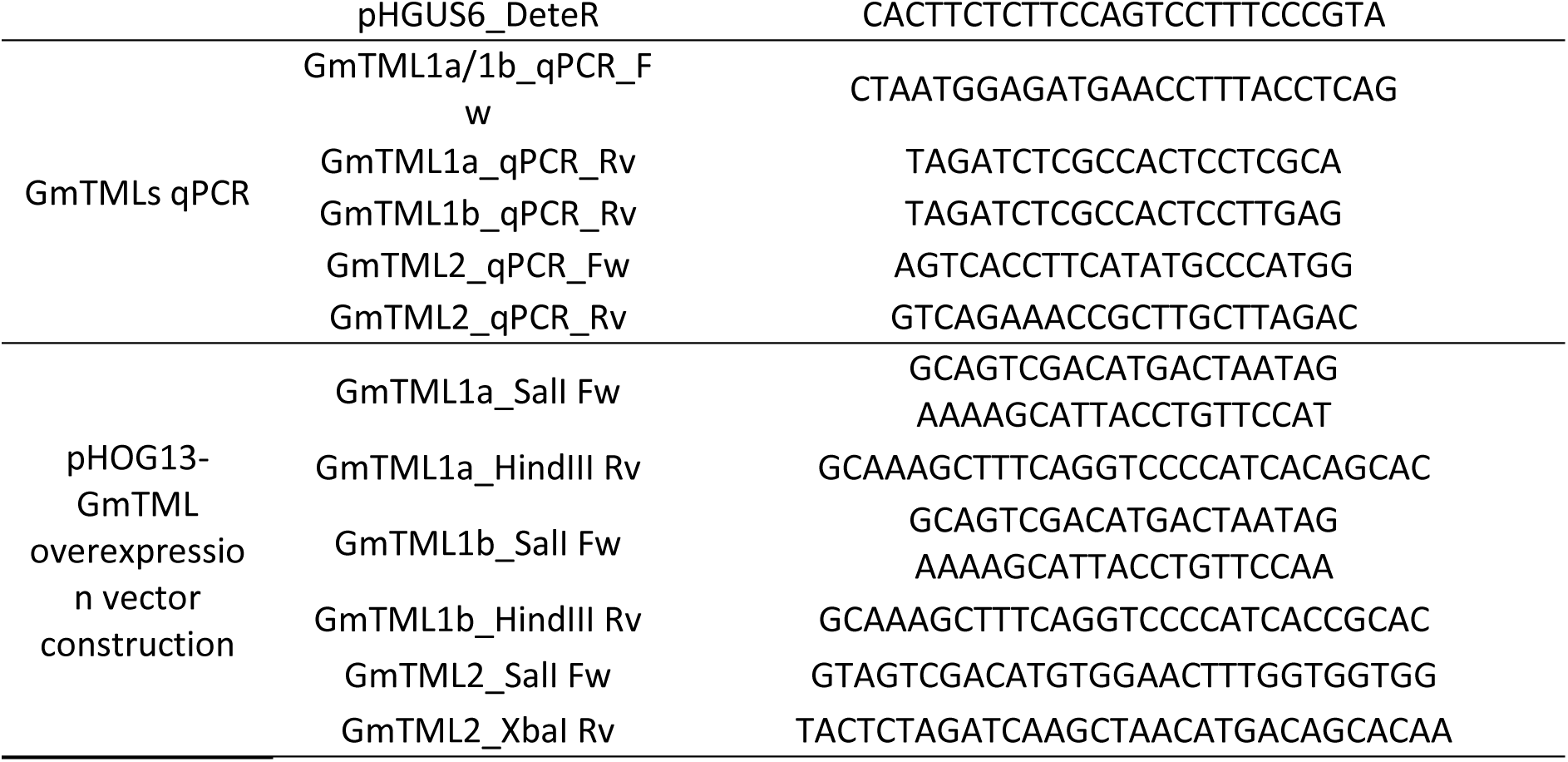
Primer sequences used for the construction and validation of pHOG13 and pHGUS7.

**Table S3.**
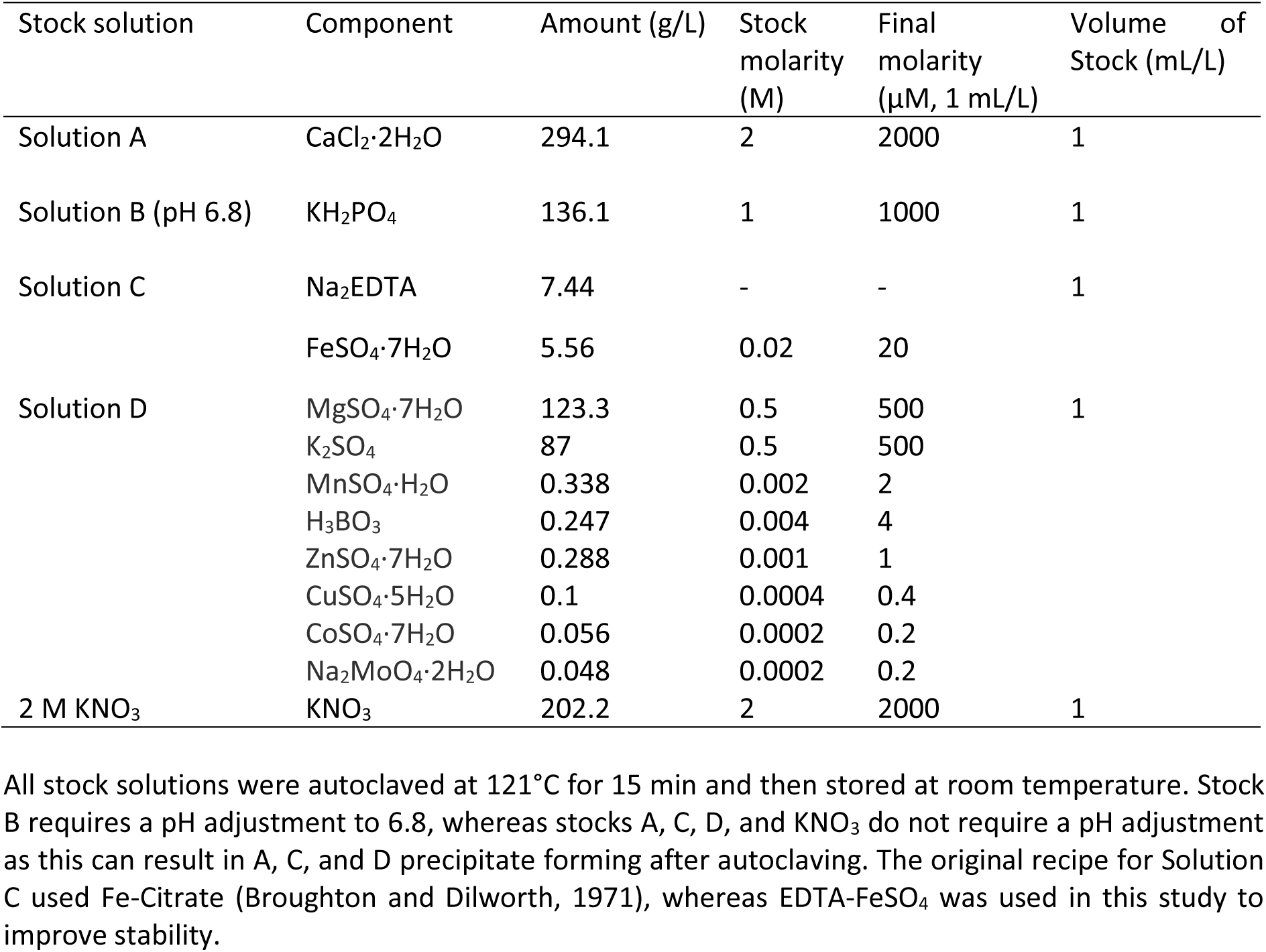
B&D and KNO_3_ nutrient solutions used to fertilise plants.

| Stock solution | Component | Amount (g/L) | Stock molarity (M) | Final molarity (μM, 1 mL/L) | Volume of Stock (mL/L) |
| --- | --- | --- | --- | --- | --- |
| Solution A | CaCl<sub>2</sub>·2H<sub>2</sub>O | 294.1 | 2 | 2000 | 1 |
| Solution B (pH 6.8) | KH<sub>2</sub>PO<sub>4</sub> | 136.1 | 1 | 1000 | 1 |
| Solution C | Na<sub>2</sub>EDTA | 7.44 | - | - | 1 |
|  | FeSO<sub>4</sub>·7H<sub>2</sub>O | 5.56 | 0.02 | 20 |  |
| Solution D | MgSO<sub>4</sub>·7H<sub>2</sub>O | 123.3 | 0.5 | 500 | 1 |
|  | K<sub>2</sub>SO<sub>4</sub> | 87 | 0.5 | 500 |  |
|  | MnSO<sub>4</sub>·H<sub>2</sub>O | 0.338 | 0.002 | 2 |  |
|  | H<sub>3</sub>BO<sub>3</sub> | 0.247 | 0.004 | 4 |  |
|  | ZnSO<sub>4</sub>·7H<sub>2</sub>O | 0.288 | 0.001 | 1 |  |
|  | CuSO<sub>4</sub>·5H<sub>2</sub>O | 0.1 | 0.0004 | 0.4 |  |
|  | CoSO<sub>4</sub>·7H<sub>2</sub>O | 0.056 | 0.0002 | 0.2 |  |
|  | Na<sub>2</sub>MoO<sub>4</sub>·2H<sub>2</sub>O | 0.048 | 0.0002 | 0.2 |  |
| 2 M KNO<sub>3</sub> | KNO<sub>3</sub> | 202.2 | 2 | 2000 | 1 |

All stock solutions were autoclaved at 121°C for 15 min and then stored at room temperature. Stock B requires a pH adjustment to 6.8, whereas stocks A, C, D, and KNO<sub>3</sub> do not require a pH adjustment as this can result in A, C, and D precipitate forming after autoclaving. The original recipe for Solution C used Fe-Citrate (Broughton and Dilworth, 1971), whereas EDTA-FeSO<sub>4</sub> was used in this study to improve stability.

**Table S4.** Minimal medium (MIN) used for culturing *Agrobacterium rhizogenes* K599.

| Reagent | Amount to add (g) |
| --- | --- |
| K<sub>2</sub>HPO<sub>4</sub> | 0.5 |
| MgSO<sub>4</sub>•7H<sub>2</sub>O | 0.2 |
| NaCl | 0.1 |
| Mannitol | 10 |
| (NH<sub>4</sub>)<sub>2</sub>SO<sub>4</sub> | 1 |
| Malic acid | 1.34 |
| Agar | 15 |
| Milli-Q H<sub>2</sub>O | up to 1 L |

**Fig. S1.**
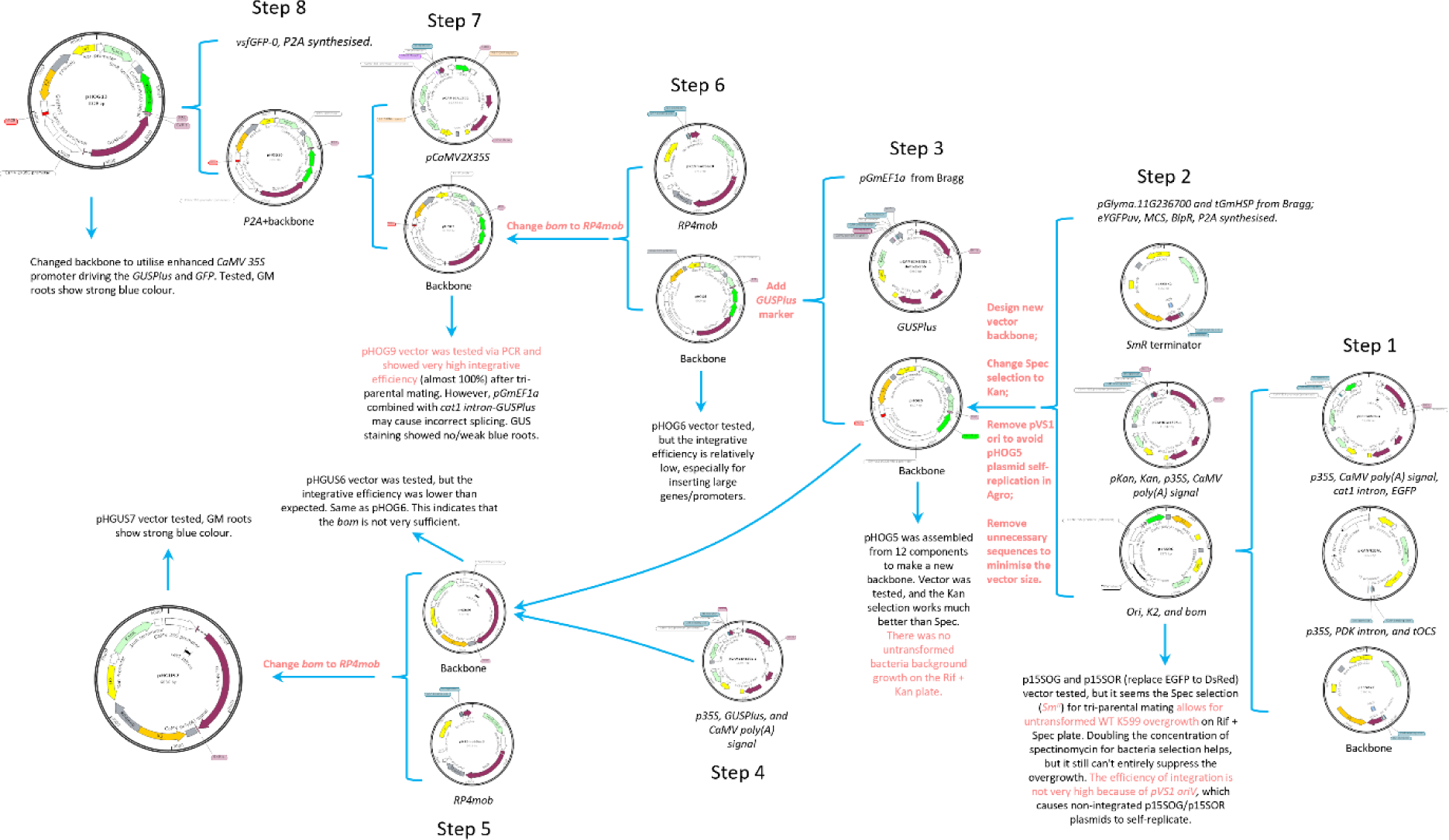
The workflow of integrative vector construction (only key intermediate vectors are shown). Step 1: p15SOG and p15SOR vector construction. Utilised p15SRK2 as an initiation vector backbone, PCR amplified *p35S, CaMV poly(A)* signal*, cat1* intron*, EGFP* from pGFPGUSplus, *DsRed* from pHairyRed, and *p35S, PDK* intron*, and tOCS* from pKANNIBAL. All the fragments were cloned into p15SRK2 through combined Gibson Assembly and In-Fusion seamless cloning strategies. Step 2: pHOG5 vector assembly. Vector backbone assembled from scratch, which is *Ori, K2,* and *bom* from p15SOG; *pKan, Kan, p35S, CaMV poly(A)* signal from pCAMBIA1305.1; *SmR* terminator from p15SRK2; *pGlyma.11G236700 and tGmHSP* from *G. max* Bragg genomic DNA*; eYGFPuv,* MCS*, BlpR, P2A* from synthesised fragments using Golden Gate Assembly and Gibson Assembly. Note: earlier versions of p15SOG and p15SOR used *Spec^R^*for selection, which caused extensive bacteria overgrowth during tri-parental mating, even when doubling the concentration of spectinomycin, and so this was swapped with *Kan^R^*. Step 3: pHOG6 vector construction. *pGmEF1a* from *G. max* Bragg and *GUSPlus* from pCAMBIA1305.1 all PCRed and cloned into the new vector backbone pHOG5. Step 4: Generated a new promoter::*GUS* integrative vector backbone pHGUS6. DNA fragments for *p35S, GUSPlus, and CaMV poly(A)* signal amplified from pCAMBIA1305.1 are cloned with minimum backbone fragments from pHOG5 using NEBuilder HiFi DNA Assembly strategy. Step 5: Final promoter::*GUS* vector pHGUS7 with a better mobility element. Replace the *mob* with *RP4mob* from pK19mobSac8 plasmid. Step 6: Modified pHOG6 to create pHOG9 using a better mobility element. PCR amplified the *RP4mob* fragment from pK19mobSac8 plasmid ligated with linearised pHOG6 backbone without *mob* element. Step 7: pHOG10 vector construction. Combine *pCaMV2X35S* with pHOG9 vector using NEBuilder HiFi DNA Assembly. Step 8: Final overexpression integrative vector pHOG13 construction. Synthesised *vsfGFP-0*, ligated with the *GUSPlus* gene, utilises *P2A* as a linker to achieve *GUSPlus* and *vsfGFP-0* gene drive under the same promoter but separated during translation. Each step after the vector construction was tested through tri-parental mating followed by hairy root transformation to verify and guide the next step’s optimisation.

## Notes

### Competing Interest Statement

The authors have declared no competing interest.

